# Ablation of GM3 Gangliosides in cardiomyocytes modestly impacts heart size but does not protect the murine heart against ischemia reperfusion injury

**DOI:** 10.64898/2026.07.13.738350

**Authors:** Yow Keat Tham, Daniel G. Donner, Gunes S. Yildiz, Helen Kiriazis, Aya Matsumoto, Kyah Grigolon, Emma I. Masterman, Natalie A. Mellett, Teleah G. Belkin, Jieting Luo, Akshima Dogra, Aascha D’Elia, Peter J. Meikle, Julie R. McMullen

## Abstract

Advances in mass spectrometry have seen the identification of hundreds of new lipid species, some of which have been found to be associated with adverse cardiac remodeling. Key among these are GM3 gangliosides, which have been associated with metabolic disease, and more recently, adverse cardiac remodeling. Whether GM3s have a direct pathophysiological effect in the heart remains unclear. The present study investigated the effects of cardiomyocyte-specific knockout of GM3 synthase (GM3S, enzyme responsible for the synthesis of GM3) in the heart under basal settings and in response to ischemia-reperfusion (I/R) injury.

A new cardiomyocyte-specific GM3S knockout (KO) model was generated, with knockout confirmed via lipidomic profiling. Under basal conditions, male GM3SKO mice exhibited reduced heart weight to tibia length (HW/TL) ratios with no evidence of pathological remodeling, while female mice showed no significant morphological differences. Male GM3SKO mice subjected to 1 hour ischemia and 4 weeks reperfusion demonstrated reduced HW/TL ratio compared to control mice subjected to I/R. However, no significant differences were observed in cardiac function, heart failure and fibrotic markers.

Lipidomic profiling (49 classes, ∼850 species) revealed significant accumulation of dihexosylceramide, a metabolic precursor of GM3 in the male heart under basal and post-I/R conditions. In male GM3SKO I/R hearts, GM3 reduction was associated with decreases in odd- and branch-chained phospholipids, together with distinct changes in circulating ether lipid species.

Collectively, cardiomyocyte-specific GM3 depletion contributed to sphingolipid remodeling but did not confer protection against I/R-mediated injury. These findings suggest that elevated GM3 levels observed in settings of cardiac pathology are not cardiomyocyte driven, highlighting the importance of understanding cell-type specific contributions to adverse cardiac remodeling.

## Introduction

Heart failure remains a major global health and economic burden, affecting up to 56 million people and accounting for $139 billion in direct expenditure in 2021 ^1,2^. With increasing cardiometabolic risk factors and an ageing population, this burden is projected to increase. While current therapies slow disease progression, they do not reverse established pathology, further highlighting the need for novel therapeutic targets.

Advances in mass spectrometry technology and methodology in the past two decades have transformed lipidomics, allowing for high throughput and/or high-resolution profiling of lipid species in health and disease ^3–5^. The application of lipidomics has uncovered new therapeutic targets and biomarkers across conditions including Alzheimer’s disease, cancer, and type 2 diabetes ^6–8^.

The pathological roles of “traditional” clinical lipids such as cholesterol and triacylglycerides in cardiovascular disease (CVD) are well established, particularly in lipid overload states such as atherosclerosis and diabetic cardiomyopathy ^9–11^. More recent lipidomic studies have demonstrated that cardiac and circulating lipidomes are altered following specific cardiac insults such as myocardial infarction, atrial fibrillation and cardiac hypertrophy ^12–16^.

GM3 gangliosides are a class of lipids derived downstream of the sphingolipid biosynthesis pathway which also produces ceramides and sphingomyelins. GM3s are lipids with important physiological roles in structure (plasma membrane rafts) and signaling, and are abundant in the liver, adipose tissues and human milk ^17–19^. Increased GM3 levels have previously been associated with a variety of metabolic diseases such as obesity and type 2 diabetes ^20–22^. Inclusion of circulating GM3 lipid species in patients with diabetes (ADVANCE patient cohort) improved the detection and prediction of atrial fibrillation ^14^. More recently, veteran athletes with atrial fibrillation demonstrated increased circulating GM3 levels compared with veteran athletes without atrial fibrillation ^23^. Cardiac GM3 levels were also elevated in cardiac tissue of mouse models of heart failure and atrial fibrillation (HF+AF), and reduced activity of a cardioprotective kinase, phosphoinositide 3-kinase activity ^24,25^. By contrast, cardiac GM3 levels were unchanged or tended to be lower in cardiac tissue of mouse models with physiological cardiac hypertrophy ^15,25^. Further, a small molecule that reduced cardiac GM3 levels was associated with better outcomes in a mouse model with HF+AF ^24^.

Collectively, these findings suggest that GM3 accumulation is a consistent feature of cardiac pathology and reducing it may be beneficial. However, whether GM3 exerts a direct pathophysiological role in the heart remains unknown. The aim of the present study was to generate a new mouse model with cardiac myocyte specific deletion of GM3 Synthase (GM3S; GM3S is the enzyme responsible for the synthesis of GM3) to investigate the pathophysiological role of GM3 lipids in cardiomyocytes, and its impact on the heart.

## Methods

### Animal care and experimentation

Animal care and experimentation were conducted following protocols approved by the Alfred Research Alliance Animal Ethics Committee, VIC, Australia (ethics #: E/1847/2018/B and E/8127/2021/B). Mice were housed in the Alfred Medical Research and Education Precinct Animal Centre under a 12-hr light-dark cycle, temperature-controlled environment. All mice were fed a standard irradiated rat and mouse feed (Specialty Feeds).

### Generation of cardiac specific GM3SKO mice

Cardiac specific GM3SKO mice were generated via the Cre/loxP system. GM3S^flox/flox^ mice on C57BL/6 background were crossed to α*MHC-Cre* mice (Cre^Tg/-^) on a FVB/N background. This resulted in the specific deletion of GM3S from the cardiomyocytes shortly after birth. This breeding strategy produced F1 progeny of ∼50% GM3S ^flox/+^Cre^Tg/-^ and GM3S ^flox/+^Cre^-/-^. These mice were cross-bred to produce six genotypes: GM3S ^flox/flox^Cre^Tg/-^ (KO), GM3S ^flox/flox^Cre^-/-^ (FC), GM3S ^flox/+^Cre^Tg/-^ (Het), GM3S ^flox/+^Cre^-/-^ (½ FC), GM3S ^+/+^Cre^Tg/-^ (Cre Ctl), GM3S ^+/+^Cre^-/-^ (WT). This strategy was utilised for the basal phenotyping cohort. Additional breeding between FC and KO mice was performed to generate only KO and FC mice only for the I/R cohort. Genotypes were determined via PCR as described previously (primer details; Supp Table 1). All mice from F1 progeny were on the same mixed genetic background. Tissue morphology comparison of all control mice (FC, ½ FC, Cre Ctl, WT) demonstrated no significant difference – subsequently all were combined as “Controls; Ctls” in the basal phenotyping cohort. The Cre transgenic mice alone have previously been shown to develop no morphological, functional or molecular aberrations.

### Experimental cohorts and animal inclusions/exclusions

#### Basal phenotyping cohort

Morphological phenotyping was performed in adult (∼ 15 weeks old) cardiac specific GM3SKO mice and littermate controls of both sexes under basal conditions. There were no exclusions (Males; Ctls = 13, Hets = 4, KO = 9; Females; Ctls = 11, Hets = 8, KO = 10).

#### Ischemia Reperfusion cohort

Adult (∼ 12 weeks old) cardiac specific GM3SKO mice and littermate controls of both sexes were subjected to ischemia reperfusion (I/R) surgical procedure. Morphological phenotyping was assessed, alongside gene expression and cardiac function analyses. (Males; Ctls = 5, KO = 7; Females; Ctls = 4, KO = 3).

### Ischemia reperfusion

Briefly, mice were anesthetized with Ketamine/Xylazine/Atropine (KXA; 100/20/1.2mg/kg, i.p.) before undergoing a surgical procedure where the left anterior descending coronary artery was reversibly ligated to induce 1 hour ischemia, followed by reperfusion as previously described ^26^. Echocardiography was also conducted 24 hours post I/R surgery for spatial assessment of the stunned left ventricular myocardium. This serves as a surrogate analysis of area-at-risk (AAR), which serves as an exclusion criterion for mice used in the study (ie. Excluding mice with AAR <35% or >55%).

### Cardiac echocardiography

Echocardiography was conducted with the Vevo 2100 Fujifilm Visual Sonics Ultrasound with a MS series 550D probe. Echocardiography was performed on mice anaesthetized on 1.8% isoflurane, with heart size and function assessed via parasternal long-axis (pLAX) B-mode ^27^.

### Tissue collection

At study endpoint, mice were anesthetized with sodium pentobarbitone (Lethabarb; 80 mg/kg, i.p.). Once mice had reached an adequate level of anesthesia (assessed by the pedal reflex test), blood was collected via cardiac puncture via a 25G needle and 1 mL syringe. This procedure was followed by cervical dislocation. Blood collected was stored on ice in K2EDTA tubes (Becton Dickinson) then centrifuged at 4°C, 4,000 g for 10 min. Plasma was collected into fresh 1.5mL tubes and stored at -80°C. Tissues and plasma were collected for molecular and lipidomic analyses. Tibias were collected to normalize for organ weights.

### RNA extractions and RT-qPCR

RNA was extracted from frozen tissues with a probe homogenizer (PRO Scientific, Model Pro 200) with TRI regent (Sigma-Aldrich) following manufacturer’s instructions. RNA was quantified using a Nanodrop Spectrometer (Thermo Fisher Scientific).

cDNA was synthesized from 2μg of total RNA via reverse transcription using High Capacity cDNA Reverse Transcription Kit (Life Technologies, Thermo Fisher Scientific). qPCR was conducted using TaqMan Gene Expression Assays with TaqMan Fast Universal Master Mix (2x). Amplification was performed on an Applied Biosystems 7500 or Quant Studio 6 or 7 Flex real-time PCR instrument. Gene expression was quantified by normalizing to hypoxanthine phosphoribosyltransferase 1 (*Hprt1*) using the 2^-ΔΔCt^ method. Details of probes used are provided in Supp Table 2.

### Lipidomics analyses

Lipidomic profiling was conducted on cardiac tissue and plasma from mice as previously described ^3,15,28^. In brief, heart tissues were homogenized in ∼100μL 1xPBS using the Branson digital probe sonicator. Protein concentration of tissue homogenates was determined using the PierceTM BCA protein assay kit. Homogenates were diluted with 1xPBS to 5mg/mL. 10μL of internal standard and 200μL of chloroform:methanol was added to each sample (10μL of heart homogenate or plasma) before being placed in a bath sonicator at room temperature for 30 minutes. Samples were then spun down at 16,060g for 10 minutes. 200μL of supernatant was pipetted into a 0.5mL polypropylene 96-well plate and dried down via a Speedivac and pump. Each sample had 50μL methanol with ammonium formate added before they were centrifuged at 4000g for 5 minutes. 100μL of each sample was then aliquoted from the 96-wells to glass vials. Lipidomic analyses were conducted via liquid chromatography via the Agilent 1290 liquid chromatography system together with the Agilent 6495C triple quadruple mass spectrometer as previously described ^28^.

### Statistical analyses

Results are presented as mean±SEM unless noted otherwise. For the basal phenotyping cohort, a one-way ANOVA followed by Tukey’s post hoc analysis was utilized. For the I/R cohort, an unpaired t-test was used to compare two groups. For lipidomics analyses, an unpaired t-test was applied, followed by a Benjamini Hochberg correction. A value of p<0.05 was considered significant. All relative units are expressed as a fold change with the relevant control group normalized to 1.

## Results

### Generation of a new mouse model with reduced GM3 lipids in the heart

As described earlier, cardiac myocyte-specific GM3SKO mice were generated using a Cre-loxP targeting approach (Fig 1A). Lipidomics was performed to confirm specific knockdown of GM3 lipids in the heart. Total GM3 lipid levels were decreased by 86.8% and 87.5% in the heart of male and female KO mice but not in heterozygote mice (Figure 1B). There was no significant difference in total GM3 in the plasma or liver (Figure 1C). Reduced GM3 in the heart was reflected in all individual GM3 species in male KO mice, and the majority of individual GM3 species in female KO mice (Figure 1D).

**Figure 1.**
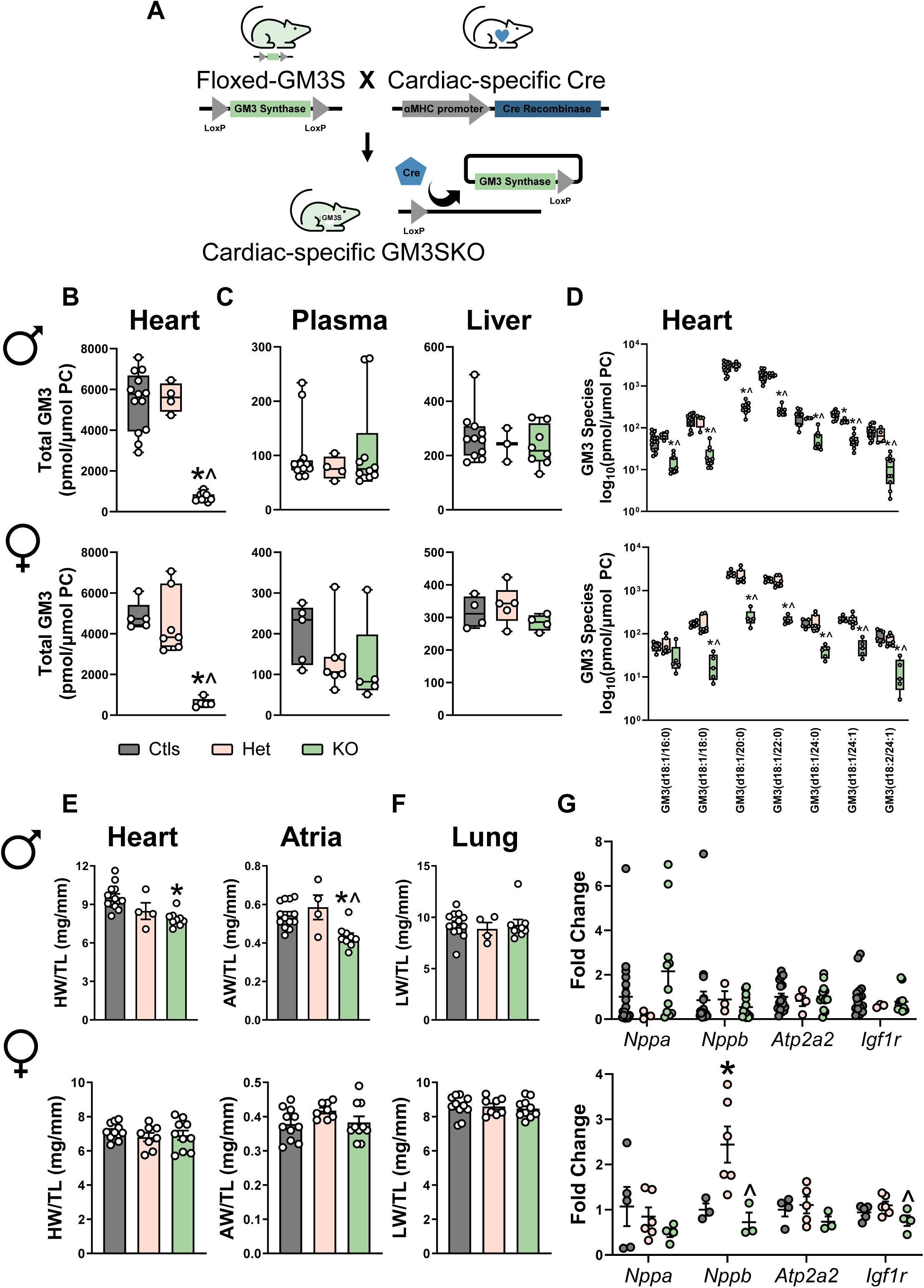
Basal phenotyping GM3SKO mice. (A) Schematic showing the generation of the new genetic mouse model. GM3S was deleted specifically from adult cardiac myocytes by breeding floxed GM3S mice with the αMHC-Cre transgenic mouse. Cardiac-specific expression of Cre excises GM3 synthase in cardiac myocytes only, leading to reduction of GM3S specifically in the heart. (B-C) Lipid analysis of total GM3 species in (B) heart, (C) plasma and liver of male and female mice. (D) Lipid analysis of individual GM3 species in hearts of male and female mice. Morphological analyses showing (E) heart and atria weight/tibia length ratios, (F) Lung weight/tibia length ratios of male and female mice. (G) Gene expression of markers of heart failure, contractility and cardiac health from hearts of male and female mice. One way ANOVA with Tukey’s post hoc were employed for statistical analyses. *p<0.05 vs Ctls, ^p<005 vs Het.

### Reduced GM3 lipids in the heart had a modest impact on heart size in male but not female mice

In male mice, body weight and tibia length were comparable between groups (Supp Table 3). Heart and atria weight/tibia length ratios (HW/TL, AW/TL) were significantly lower in male GM3SKO mice vs controls (Figure 1E). Female GM3SKO mice were modestly heavier than controls, but there was no difference in tibial length (Supp Table 3). Unlike the male mice, there were no significant differences in HW/TL or AW/TL in female GM3SKO mice. Normalized lung wet weights, which can increase in mice with a failing heart due to fluid accumulation in the lung were not significantly different between GM3SKO mice vs controls of both sexes (Figure 1F). Next, we assessed cardiac gene expression of molecular markers of cardiac stress (ANP-*Nppa*, BNP-*Nppb*) contractility (Serca2a-*Atp2a2*) and the IGF1 pathway (Igf1r, IGF1 pathway that can be regulated by GM3 (28, 29)). These markers were largely unchanged between GM3SKO and control/heterozygote mice, though BNP was elevated in the ventricle of heterozygote female mice vs control and Igf1r expression was lower in GM3SKO vs heterozygote female mice (Figure 1G).

### GM3SKO mice are not significantly protected against cardiac I/R injury

Previous studies conducted in our laboratory demonstrated that increased GM3 levels in the heart were associated with cardiac dysfunction and pathology ^15,24,25^. Despite a limited phenotype in GM3SKO males and no obvious phenotype in GM3SKO females under basal conditions, there are numerous examples of pathological or protective phenotypes only becoming apparent under cardiac stress settings ^29–34^. Given that, male and female GM3SKO and FC mice were subjected to 1-hour cardiac I/R injury, with tissues collected at 28 days post-surgery. Post-I/R, body weights of male GM3SKO mice and floxed controls (FCs) were not different (Figure 2A, Supp Table 4). Area at risk assessed 24h post-surgery showed no differences between genotypes, indicating consistency in degree of injury from I/R surgery (Figure 2B). HW/TL ratio of GM3SKO mice were significantly lower following I/R vs FCs (Figure 2C, Supp Table 4), and this was attributed to a significant decrease in only the left ventricular chamber (Figure 2D, Supp Table 4). Additionally, there was a trend for a reduction in the right normalized atrial weight, but this trend was not observed in the left atrial weight, or indeed the combined atrial weights (Figure 2E, Supp Table 4). Lung weight/ tibia length ratio of GM3SKO mice was unchanged vs FCs following I/R injury (Figure 2F, Supp Table 4). In addition, at 28 days post I/R there were no differences in cardiac function via echocardiography based on ejection fraction at comparable heart rates assessed via echocardiography (Figure 2G, H, Supp Table 5). Gene expression assessment of molecular markers of heart failure and fibrosis were not significantly different between genotypes after IR injury, although the contractility marker *Atp2a2* (Serca2a) was significantly lower in GM3SKO mice (Figure 2I). No significant differences were identified in female mice subjected to I/R based on morphology, cardiac function or molecular makers (Supp Figure 2, Supp table 5).

**Figure 2.**
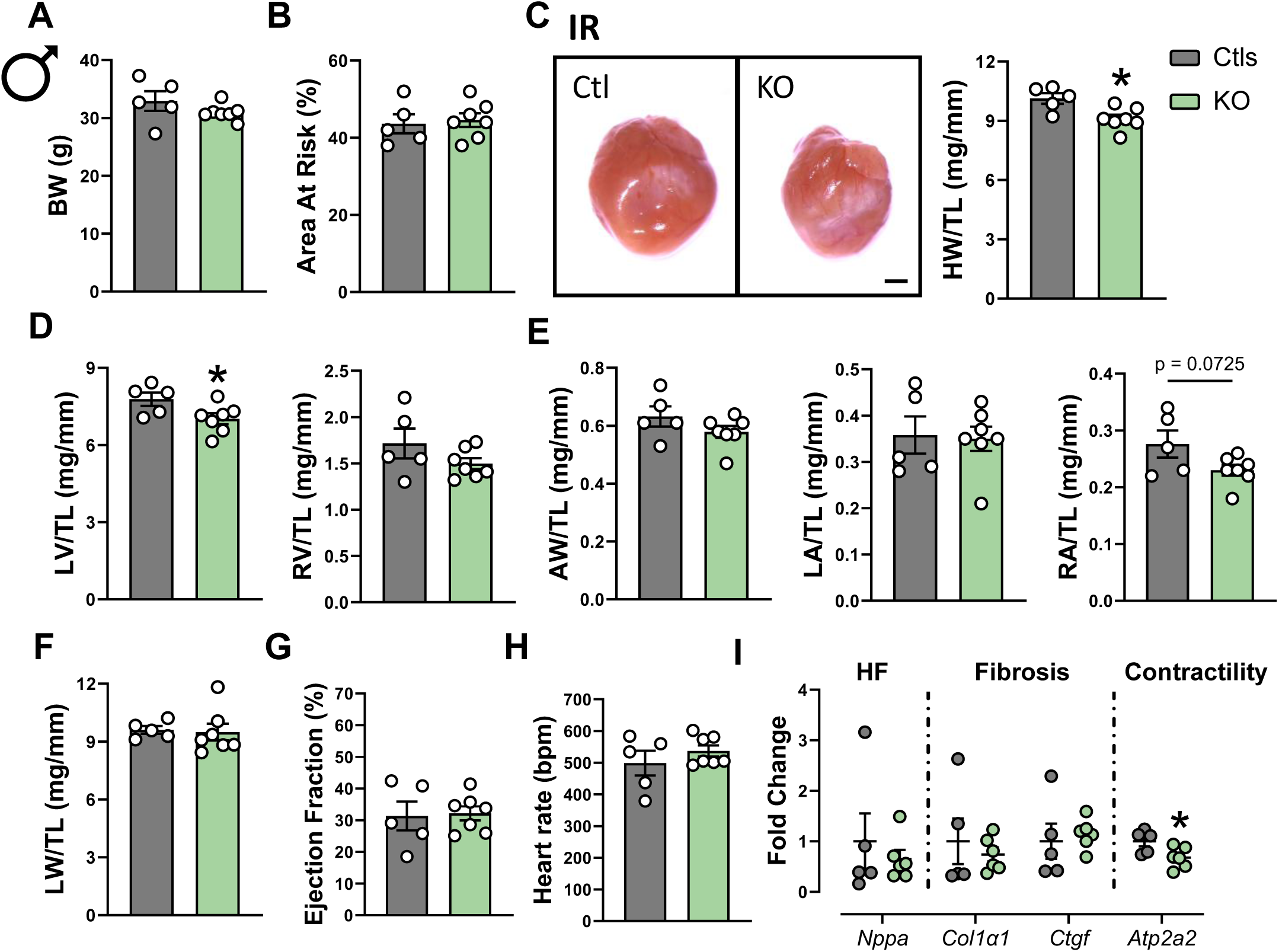
Phenotyping male GM3SKO mice following I/R injury. Scatter plots demonstrating (A) body weight and (B) area at risk assessed 24 hours after I/R surgery. (C) Representative images of GM3SKO hearts 4 weeks post I/R, with accompanying morphological analyses showing heart weight normalized to tibia length. Morphological analyses showing (D) left and right ventricular weights, (E) atria, with left vs right atria weights, and (F) lung weight normalized to tibia length. (G) Ejection fraction as assessed via echocardiography 4 weeks post I/R. (H) Gene expression of markers of heart failure, fibrosis and contractility from hearts of mice 4 weeks post I/R. Unpaired t-tests were employed for statistical analyses. *p<0.05 vs Ctls I/R.

### Remodeling of the cardiac and circulating lipidome in GM3SKO mice under basal conditions and in response to cardiac ischemia reperfusion injury

In previous work targeting plasmalogen lipids, we identified compensatory regulation of other lipid species within the lipidome ^25^. To explore whether the absence of more distinct phenotypes under basal or I/R conditions could be due to compensatory changes to lipid species besides GM3, we interrogated the full lipid profile consisting of 49 classes and ∼850 individual lipid species between GM3SKO and control mice. GM3 lipids are synthesized via the sphingolipid biosynthesis pathway (Figure 3A). Under basal conditions, the reduction of cardiac GM3 lipids in the heart of male and female GM3SKO mice was accompanied with a 16-fold increase in total dihexosylceramide (Hex2Cer), a lipid class that precedes GM3s within the biosynthesis pathway (Figure 3B). This increase was similarly reflected in all the individual Hex2Cer lipids in GM3SKO mice of both sexes (Supp Figure 1). Notably, while no changes were observed in circulating GM3 lipids, there was a non-significant increase in circulating Hex2Cer lipids in male GM3SKO mice, with a similar pattern in females (Figure 3B, C).

**Figure 3.**
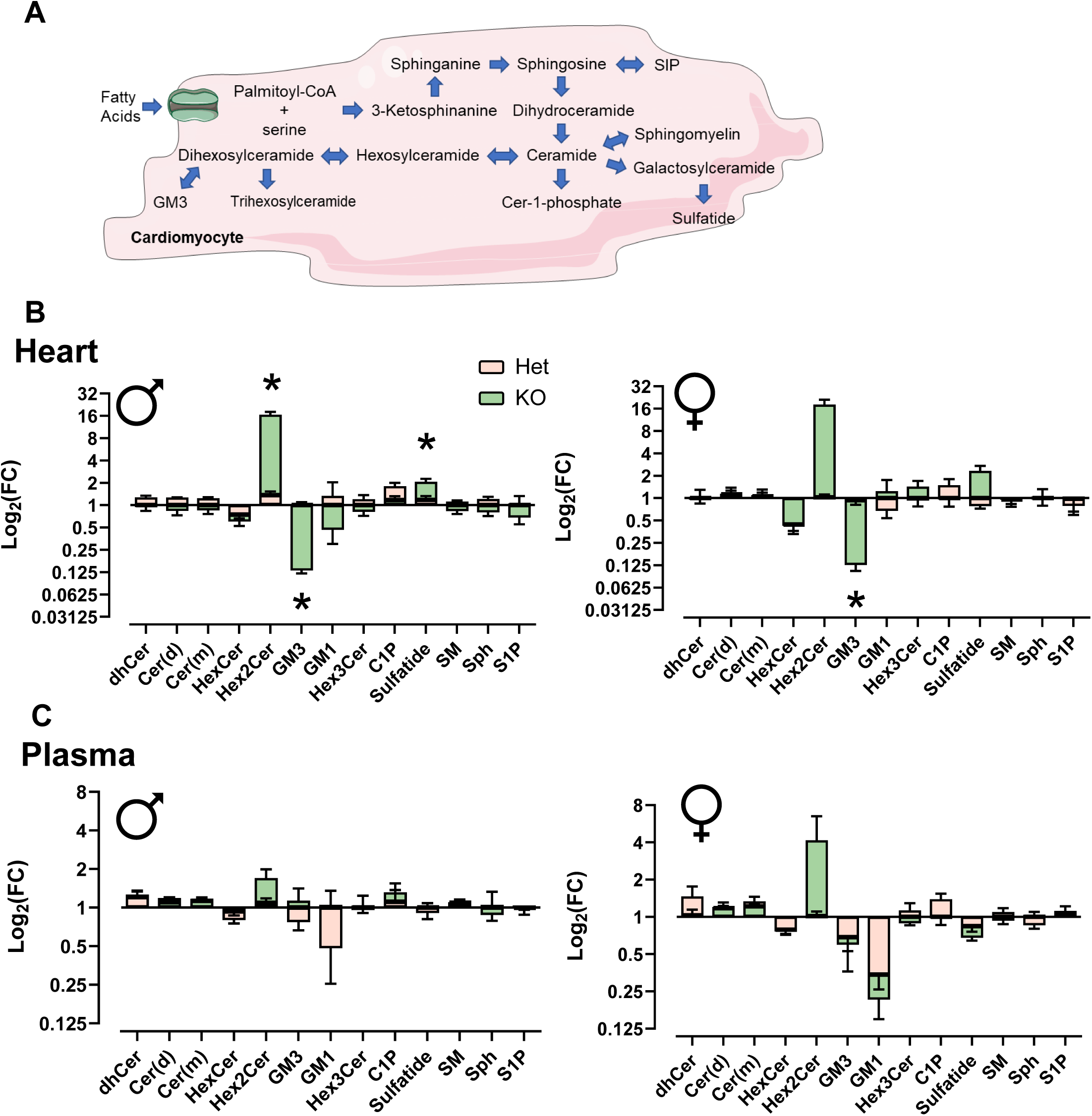
(A) Representative image of the sphingolipid biosynthesis pathway. Lipid classes within the sphingolipid domain as detected in the (B) heart and (C) plasma of male and female GM3SKO mice presented as a fold change against controls. Unpaired t-tests with Benjamini Hochberg corrections were employed for statistical analyses. *p<0.05 vs Ctls.

In the I/R model, lipidomics was performed in male hearts only because there were more non-infarcted LV mass available vs females (due to larger overall heart mass) and the animal numbers were higher. Lipidomic profiling of the GM3SKO LV subjected to I/R vs FC LV highlighted a major reduction of GM3 levels, with a concurrent increase in Hex2Cer individual lipids (Figure 4A), in line with changes observed in the cardiac lipidome under basal conditions (Figure 1A, 3B). In addition, lipid species with odd and branch chains (15-MHDA) were consistently lower in GM3SKO I/R hearts vs FCs, with these lipid species spanning across multiple phospholipid classes [LPC, PC, PE, PC(P), PE(P), PI] significantly reduced (Figure 4A).

**Figure 4.**
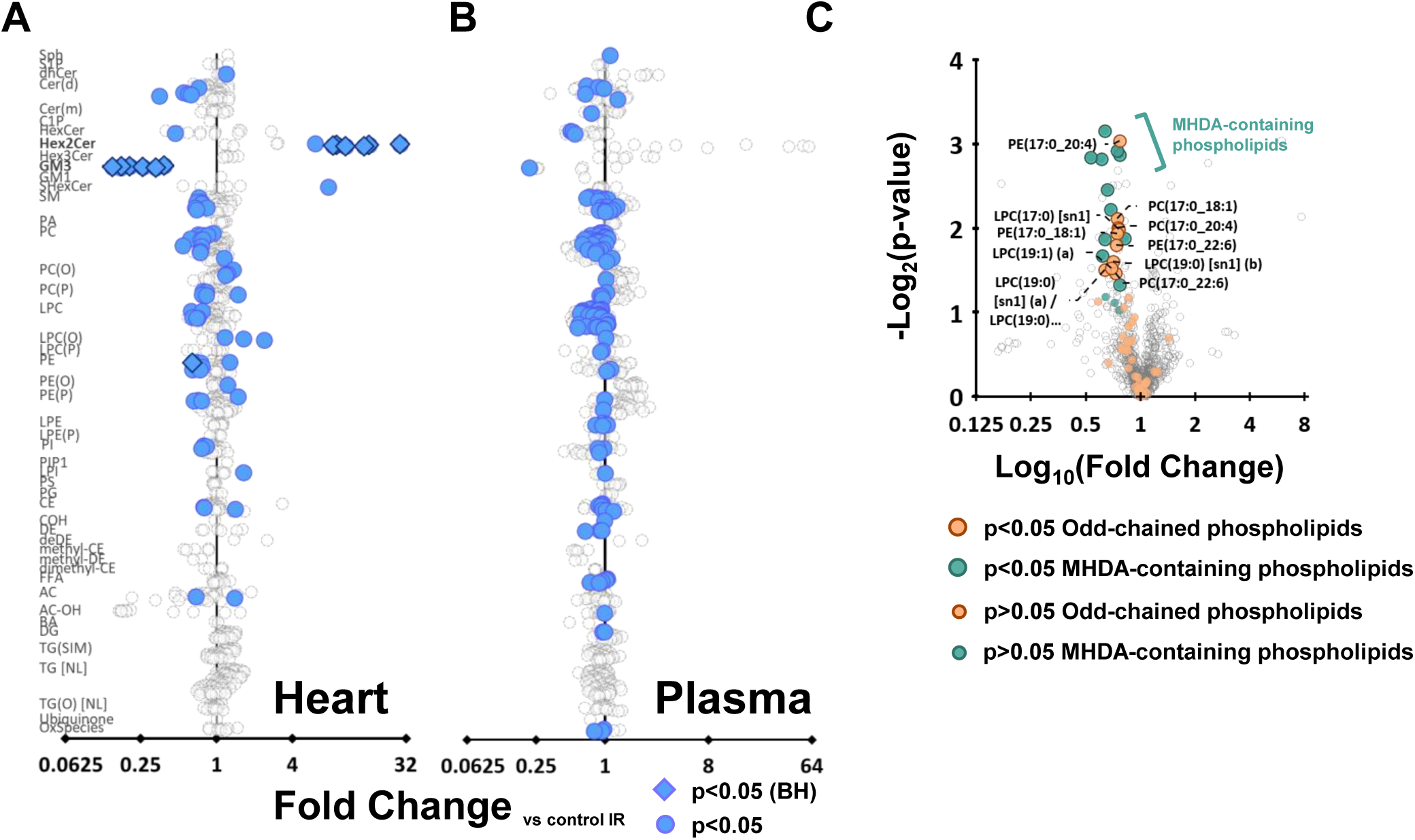
Forest plots showing differences in individual lipid species in (A) heart and (B) plasma of GM3SKO mice subjected to IR vs Ctl I/R. (C) Volcano plot highlighting change in odd-chained and MHDA-containing phospholipids in the heart of GM3SKO mice subjected to IR vs Ctl I/R. Unpaired t-tests with Benjamini Hochberg corrections were employed for statistical analyses for (A) and (B), with unpaired t-tests applied for (C). *p<0.05 vs Ctl I/R.

The circulating lipidome profiled demonstrated a comparable pattern which was not significant i.e. decrease in GM3 and increase in Hex2Cer lipid levels in GM3SKO vs FC I/R mice (Figure 4B). This trend was again similar to that observed under basal conditions (Figure 3C). There was a general trend for LPC and PC individual species to be reduced, but this was not significantly different after Benjamini-Hochberg correction (Figure 4B).

### Expression of GM3 in different cardiac cell types

The heart is a multicellular organ that consists of different cell types, including cardiomyocytes, endothelial cells, fibroblasts and immune cells. While our findings have previously associated an increase in cardiac GM3 levels with pathology, these have all been conducted in whole heart homogenates. To shed light on the potential importance of GM3 lipids in a cell type specific manner in the heart, we screened for ST3GAL5, the corresponding gene for GM3S, using the Genotype-Tissue Expression (GTEx) Portal. This revealed that ST3GAL5 expression was highest in immune, endothelial and fibroblasts but comparatively lower in cardiomyocytes (Fig 5).

**Figure 5.**
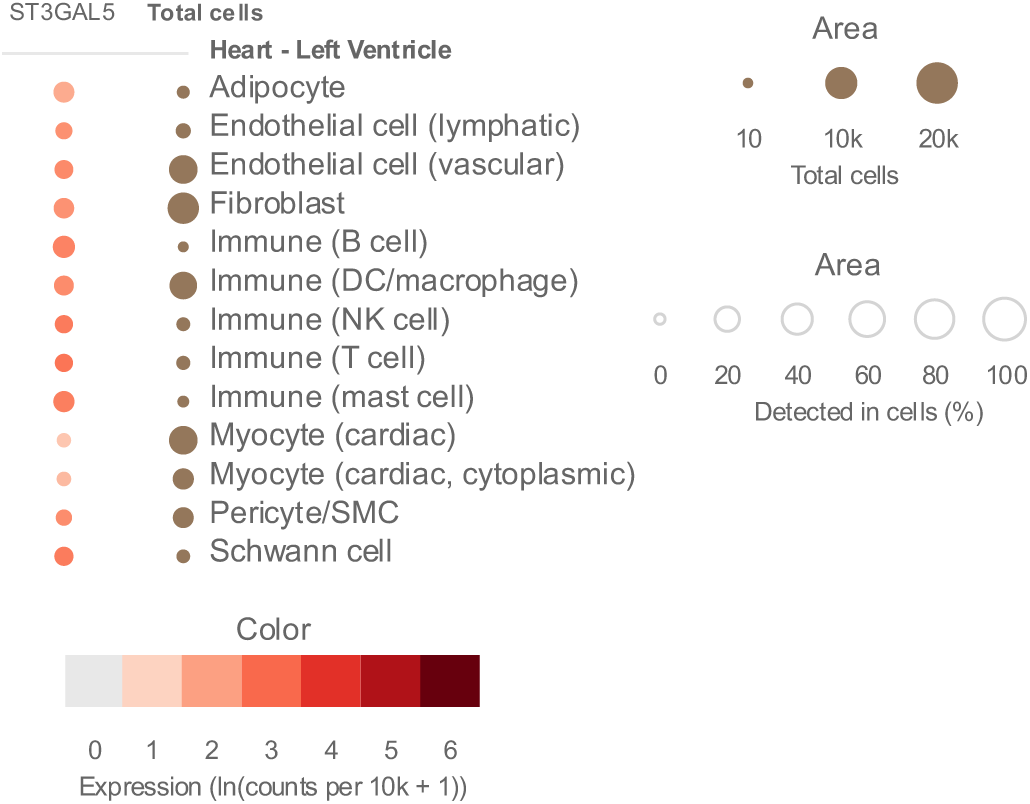
GTEx human single cell data analysis of ST3GAL5 gene expression in cell populations from the heart

## Discussion

Lipidomic profiling has increasingly been used to uncover new associations between lipid species and cardiovascular health and disease. This has enabled the identification of new therapeutic targets and/or potential biomarkers. GM3 gangliosides are a class of glycosphingolipids most abundant in the nervous system, particularly the brain, but also present in the heart ^35^. In both the rodent and human heart, GM3 represents ≥50% of all gangliosides ^36,37^. Previous studies have demonstrated a positive association between plasma GM3 levels and impaired insulin sensitivity and metabolic disease ^21,38–40^. Furthermore, global inhibition of GM3S in mouse models was associated with improved insulin sensitivity and conferred resistance to diet-induced obesity ^41,42^. More recently, our studies demonstrated similar positive associations in both cardiac and/or plasma GM3 levels with cardiac pathology ^15,24,25^. Downregulation of GM3S was also associated with reduced fibrotic markers in TGF-β activated human cardiac fibroblasts ^43^. The aim of the present study was therefore to assess whether reducing GM3 levels via cardiomyocyte-specific GM3S inhibition impacts the heart under basal conditions or in a setting of cardiac stress (I/R injury). The main findings to come from this work include 1) the generation of a new mouse model with substantial reduction of GM3 levels in the adult male and female mouse heart (>85%), 2) reduced HW/TL ratio in male GM3SKO mice under basal conditions but not females, 3) reduced normalized heart mass following I/R in males associated with lower Serca2a gene expression but with no effect on heart function, 4) a compensatory accumulation of Hex2Cer in the KO heart, a metabolic precursor of GM3, and 5) a significant dysregulation in odd and branch chained phospholipids in the hearts of male GM3SKO mice subjected to I/R injury.

### Effect of cardiomyocyte-specific GM3S knockout under basal and pathological conditions

Previous studies reported that global GM3SKO mice displayed no overall major abnormalities under basal conditions, apart from deafness, impaired activation of CD4^+^ T cells and altered cholesterol uptake ^44–47^. However, cardiac morphology and function were not specifically assessed. The tissue specific role of GM3 lipids has not been widely interrogated. Notably, Zhang et al generated smooth muscle cell-specific GM3SKO x ApoEKO mice and demonstrated that GM3 depletion enhanced ferroptosis of smooth muscle cells ^48^. To our knowledge, this is the first study to generate and study cardiomyocyte specific GM3SKO mice under basal and cardiac stress settings.

Basal phenotyping revealed that cardiomyocyte-specific GM3SKO resulted in reduced normalized heart size in male mice only. This reduction was not associated with pathological remodeling, as normalized lung weight and expression of heart failure markers were unchanged. A similar reduction in heart weight was observed in male GM3SKO following I/R injury. The response of the GM3SKO to I/R was not markedly different from control mice subjected to I/R, but cardiac Serc2a gene expression was lower in the GM3SKO. In contrast, no significant morphological differences were identified in female GM3SKO mice under basal conditions or in response to I/R injury. While this is the first instance where sexual dimorphism in GM3S deficiency has been associated with different cardiac phenotypes, a study using global GM3SKO mice reported differing levels of ADHD-like behavior between male and female mice. Additionally, they also reported an increase in body weight and reduction in both insulin receptor isoforms in peripheral tissues in male mice, while no body weight difference was observed in female mice. Conversely, both insulin receptor isoforms were unchanged in the spleen and cortex, but significantly increased in female GM3SKO livers ^49^.

The smaller hearts observed in the GM3SKO mice was an interesting phenotype as this was not accompanied with overt development of pathology. In addition, GM3SKO mice subjected to cardiac I/R injury did not demonstrate significant protection or exacerbation of pathology over FC mice subjected to I/R. This is in contrast to previous findings in transgenic mice with impaired PI3K activity in the heart, which also displayed smaller hearts with preserved heart function, but with significantly exacerbated pathology after being subjected to cardiac injury^50–53^.

Lipidomic analysis of GM3SKO hearts following I/R injury showed consistently lower levels of odd- and branched-chain (15-MHDA) lipid species (Figure 4C) compared to control hearts. A reduction in odd-chain phosphatidylcholines has been associated with impaired mitochondrial branched-chain amino acid (BCAA) catabolism in the liver ^54^. Dysregulation of BCAA catabolism may subsequently influence branched-chain phospholipid synthesis as isoleucine and leucine serve as precursors for these lipids. To our knowledge, the reduction in odd and branched chain PCs in the setting of cardiac injury has not yet been reported. Impaired BCAA catabolic flux is recognized as a feature of heart failure and cardiac stress, where interventions that either reduce excessive BCAA substrate load or enhance BCAA catabolism were associated with improved remodeling ^55,56^.

Cardiomyocyte-specific GM3S deletion resulted in a substantial accumulation of its metabolic precursor, Hex2Cer in male hearts. To our knowledge, this is the first report demonstrating Hex2Cer accumulation following GM3 knockdown in cardiac tissue. Hex2Cer, a glycosphingolipid, is known to be involved in cell proliferation, migration and angiogenesis in the context of cancer ^57,58^. Increased renal Hex2Cer levels in type 2 diabetic mouse models were associated with renal fibrosis and hypertrophy ^59^. Moreover, both surgical and phenylephrine-induced cardiac hypertrophy in mice and cell cultures were associated with elevated Hex2Cer levels, while pharmacological inhibition of Hex2Cer synthesis prevented development of hypertrophy ^60,61^. In contrast, our present study observed a decrease in heart size in males but not females, but there was no evidence of any cardiac pathology.

The relationship between GM3 lipids and metabolism remains complex. Global GM3SKO mice have demonstrated improved insulin sensitivity, enhanced wound healing response, and protection against high fat diet-induced insulin resistance (20, 29, 38). Bharathi et al further reported increased whole-body respiration and glucose oxidation in a separate global GM3SKO mouse model (55). Tissue and cell type specific studies of GM3S remain limited. Fibroblasts from the muscle of a GM3S deficient patient showed reduced oxygen consumption and mitochondrial membrane potential, and liver biopsy samples demonstrated decreased mitochondrial respiratory chain complex activity (56). In contrast, smooth muscle cell specific GM3S deletion exacerbated abdominal aortic aneurysm development, which was reversed by GM3 supplementation (45). These publications highlight how GM3S and GM3 lipids do have an important role in specific tissue or cell types. The *in silico* assessment in Figure 5 showing relatively low ST3GAL5 expression in cardiomyocytes may provide indications as to why our cardiac-specific GM3sKO demonstrated no beneficial effects in settings of cardiac injury.

## Conclusion

In the current study, cardiomyocyte-specific deletion of GM3S effectively reduced GM3s levels but did not protect against cardiac I/R injury. These findings indicate that reducing cardiomyocyte GM3 alone is insufficient to attenuate acute ischemic injury, despite previous associations linking elevated cardiac GM3 levels with pathological remodeling. Broadly, our data supports a context and cell type-specific role for GM3 in cardiovascular disease. Future studies assessing the role of GM3 lipids in non-myocyte cardiac populations will be important to fully define its role in cardiac injury and remodeling.

## CRediT authorship contribution statement

**Yow Keat Tham:** Conceptualization, Methodology, Investigation, Formal analysis, Data curation, Validation, Visualization, Writing – original draft, Writing – review & editing, Supervision, Project administration. **Daniel Donner:** Methodology, Investigation, Formal analysis, Data curation, Validation. **Gunes S. Yildiz:** Investigation, Formal analysis, Data curation, Validation. **Helen Kiriazis:** Investigation, Formal analysis, Validation. **Aya Matsumoto:** Investigation, Formal analysis. **Kyah Grigolon:** Formal analysis, Validation. **Emma Masterman:** Investigation, Formal analysis, Validation. **Natalie Mellett:** Investigation, Formal analysis, Validation. **Teleah G. Belkin:** Investigation, Formal analysis. **Jieting Luo:** Investigation, Formal analysis. **Aascha D’Elia:** Investigation. **Peter J. Meikle:** Methodology, Resources. **Julie R. McMullen:** Conceptualization, Methodology, Writing – review & editing, Supervision, Project administration, Funding acquisition, Resources.

## Acknowledgements

The Genotype-Tissue Expression (GTEx) Project was supported by the Common Fund of the Office of the Director of the National Institutes of Health, and by NCI, NHGRI, NHLBI, NIDA, NIMH, and NINDS. The data used for the analyses described in this manuscript were obtained from the GTEx Multi-Gene Single Cell Query on 02/06/2026.

---

**Supplementary Figure 1.** Forest plots of individual lipids within the sphingolipid domain in the heart and plasma of GM3SKO male and female mice vs controls. Unpaired t-tests with Benjamini Hochberg corrections were employed for statistical analyses.

**Supplementary Figure 2.** Phenotyping female GM3SKO mice following I/R injury. Scatter plots demonstrating (A) body weight and (B) area at risk assessed 24 hours after ischemia reperfusion surgery. (C) Representative images of GM3sKO hearts 4 weeks I/R, with accompanying morphological analyses showing heart weight normalized to tibia length. Morphological analyses showing (D) left and right ventricular weights, (E) atria, with left vs right atria weights, and (F) lung weight normalized to tibia length. (G) Ejection fraction as assessed via echocardiography 4 weeks post I/R. (H) Gene expression of markers of heart failure, fibrosis and contractility from hearts of mice 4 weeks post I/R. Unpaired t-tests were employed for statistical analyses. *p<0.05 vs Ctls I/R.

