## Supplementary figures and images for "Ablation of GM3 Gangliosides in cardiomyocytes modestly impacts heart size but does not protect the murine heart against ischemia reperfusion injury"

### Supplementary Figures

#### Supplementary Figure 1

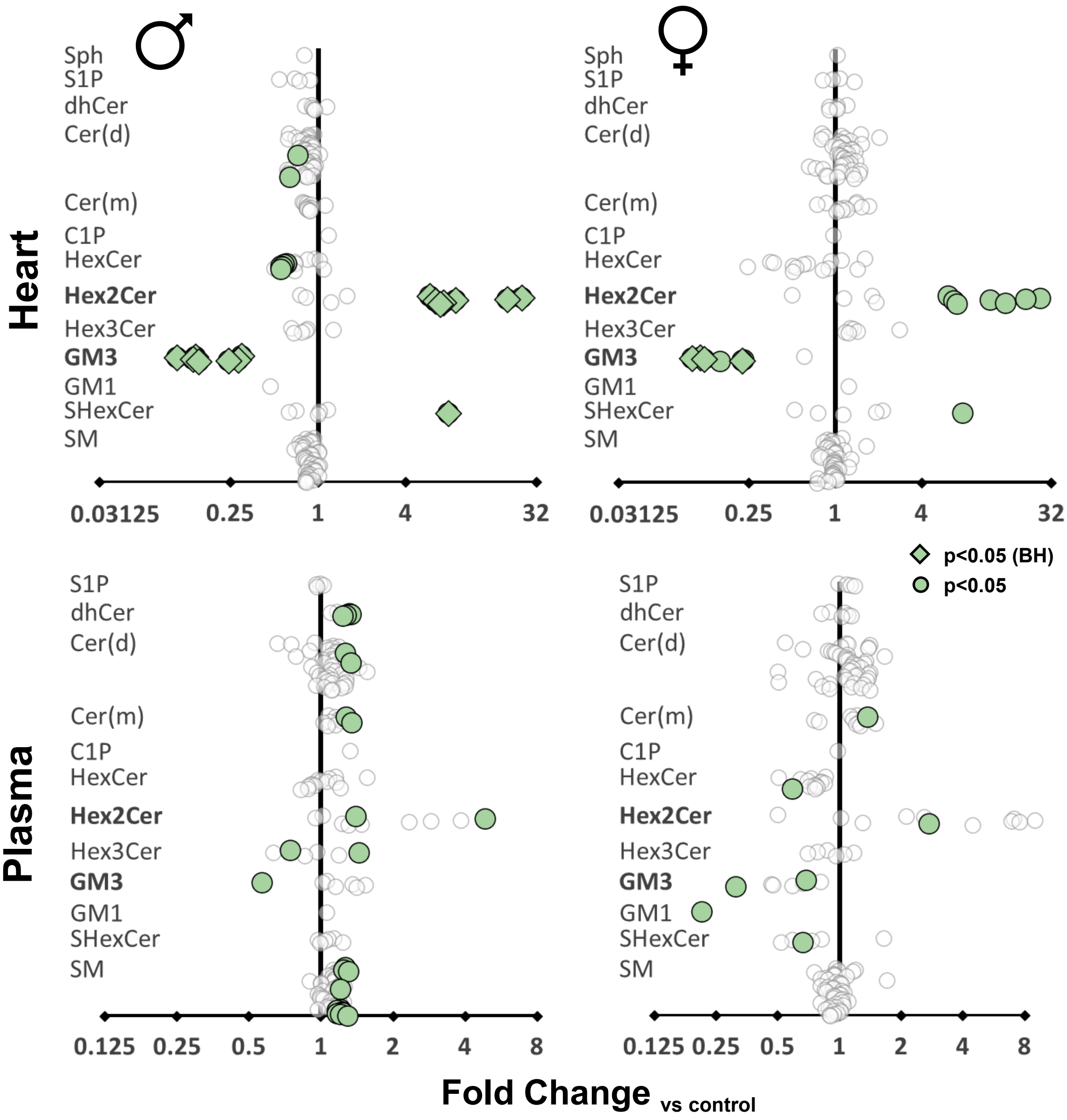

##### Supplementary Figure 2

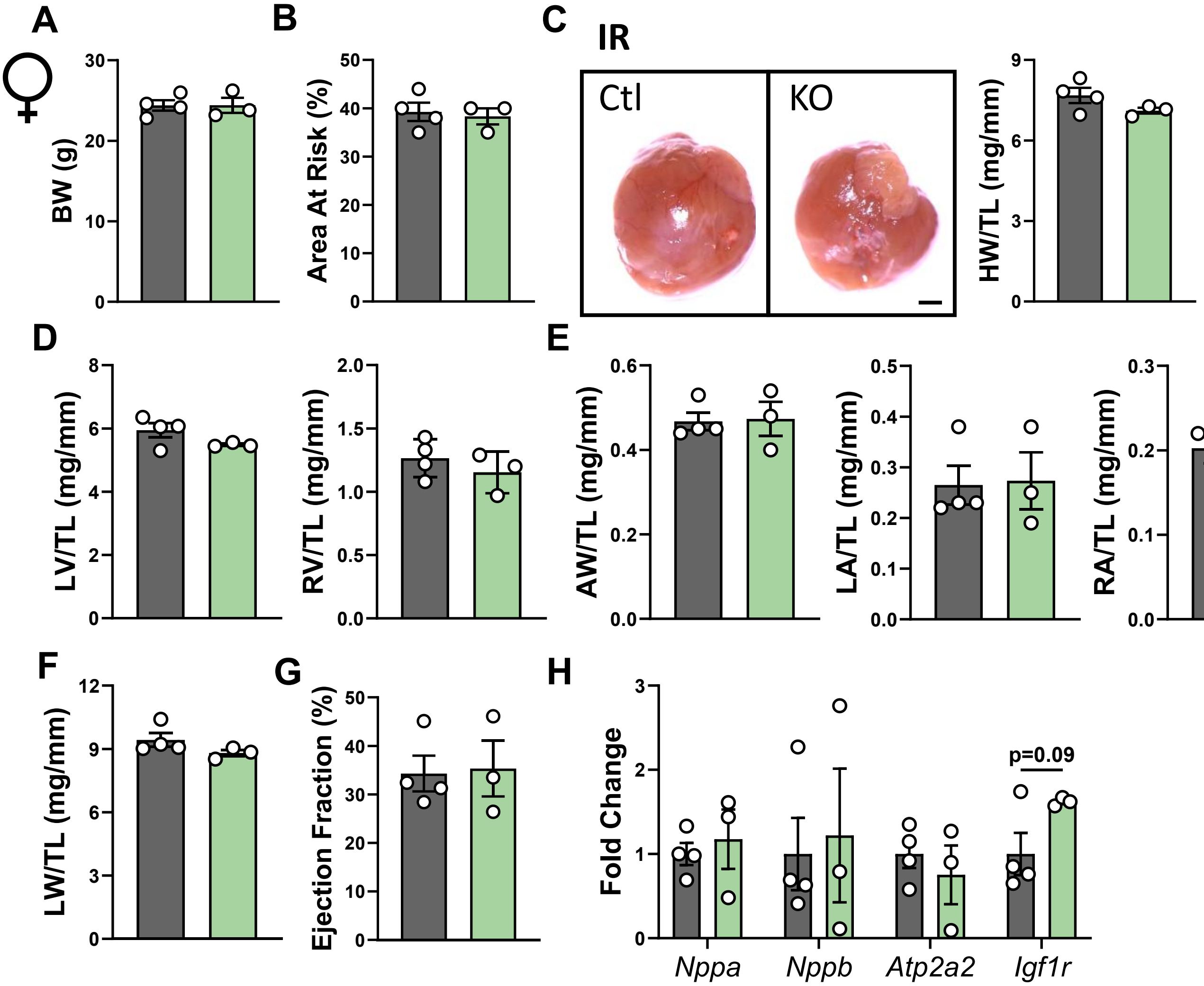
